# A STAT3-regulated lncRNA integrates microRNA biogenesis and sequestration to safeguard naïve pluripotency

**DOI:** 10.1101/2025.09.23.677977

**Authors:** Vincent Giudice, Florence Perold, Yannicke Pijoff, Nathalie Doerflinger, Nicolas Allègre, Claire Chazaud, Irène Aksoy, Pierre Savatier, Pierre-Yves Bourillot

**Affiliations:** Univ Lyon, Université Lyon 1, INSERM, Stem Cell and Brain Research Institute U1208, INRAE USC 1361, F-69500 Bron, France; Université Clermont Auvergne, CNRS, INSERM, GReD Institute, Faculté de Médecine, F-63000, Clermont-Ferrand, France

## Abstract

Leukemia inhibitory factor (LIF)/STAT3 signaling is central to maintaining naïve pluripotency in mouse embryonic stem cells (mESCs). We identify *Asgard*, a previously uncharacterized long non-coding RNA, as a direct STAT3 target required for efficient self-renewal. *Asgard* is rapidly induced by LIF, enriched in the epiblast, and its depletion reduces alkaline phosphatase–positive colony formation while enhancing differentiation. Mechanistically, *Asgard* fulfils a dual role: it acts as the primary transcript for the differentiation-promoting microRNA *Odin*, while also functioning as a sponge to sequester *Odin* and related miRNAs. This dual mechanism enables *Asgard* to both generate and buffer pro-differentiation signals, thereby stabilizing the pluripotent state while preserving responsiveness to lineage cues. Our work reveals a new paradigm in RNA-mediated control of stem cell identity, where a single STAT3-regulated lncRNA couples microRNA production with competitive inhibition to safeguard naïve pluripotency.

## Introduction

Mouse embryonic stem cells (mESCs) provide a robust system to dissect the molecular logic of naïve pluripotency. Their self-renewal is maintained by cytokine signaling through the leukemia inhibitory factor (LIF)/STAT3 pathway, which activates a transcriptional program centered on key regulators such as *Klf4*, *Tfcp2l1*, and *Esrrb* (Aksoy *et al*, 2014; Martello *et al*, 2013; Matsuda *et al*, 1999; Niwa *et al*, 2009). Constitutive activation of STAT3 is sufficient to sustain naïve pluripotency, underscoring its role as a master transcriptional regulator (van Oosten *et al*, 2012). In addition to transcription factors, non-coding RNAs (ncRNAs) have emerged as pivotal regulators of stem cell fate. Among these, microRNAs (miRNAs) form integral components of the pluripotency network (Wang *et al*, 2007). For example, the *miR-290–295* cluster reinforces self-renewal by targeting cell-cycle inhibitors and differentiation cues, while *let-7* family members promote exit from pluripotency and lineage commitment (Judson *et al*, 2009; Melton *et al*, 2010). The dynamic balance between pluripotency-supporting and differentiation-promoting miRNAs exemplifies how small RNAs act as molecular switches controlling stem cell identity.

Long non-coding RNAs (lncRNAs) add further complexity by modulating miRNA networks through multiple mechanisms (Dinger *et al*, 2009; Guttman & Rinn, 2012; Statello *et al*, 2021). First, several lncRNAs serve as primary transcripts for miRNA biogenesis, directly producing small RNAs that influence stem cell fate. Second, lncRNAs can act as competing endogenous RNAs (ceRNAs), sequestering miRNAs away from their mRNA targets and thus sustaining pluripotency gene expression (Cesana *et al*, 2011; Tay *et al*, 2014). Several ceRNAs have been directly implicated in pluripotency control. For instance, the STAT3-regulated lncRNA *Lncenc1* promotes self-renewal by sponging miR-128, thereby stabilizing *Klf4* (Monteleone *et al*, 2024). In mESCs, the lncRNA *Cyrano* sustains pluripotency by titrating miR-7, thereby preventing repression of self-renewal pathways (Smith *et al*, 2017). In human ESCs, a regulatory feedback loop involves the Large Intergenic Non-Coding RNA – Regulator of Reprogramming (*LINC-ROR*), the microRNA miR-145, and the pluripotency transcription factors *OCT4*, *SOX2*, and *NANOG*: miR-145 inhibits *OCT4*, *SOX2*, and *NANOG* expression; in turn, these transcription factors promote *LINC-ROR* transcription; and the sponge domain of *LINC-ROR* titrates miR-145, preventing degradation of pluripotency transcripts (Loewer *et al*, 2010; Wang *et al*, 2013). Another example is the sponge lncRNA *Growth Arrest Specific transcript 5* (*GAS5*), which regulates NODAL signaling in hESCs. NODAL promotes self-renewal by activating *OCT4*, *SOX2*, and *NANOG*, and *GAS5* expression is induced by *OCT4* and *SOX2*. In turn, *GAS5* sequesters miR-2467-5p, miR-3200-3p, and miR-let7a/e-5p, all of which target NODAL, thereby reinforcing pluripotency (Xu *et al*, 2016). Together, these observations suggest that lncRNAs and miRNAs form intertwined circuits that buffer pluripotency against stochastic fluctuations while preserving responsiveness to differentiation cues. However, the repertoire of STAT3-regulated lncRNAs integrating both miRNA precursor and sponge activities remains poorly characterized.

Here, we identify a novel STAT3 target lncRNA, *Asgard* (*Another Self-renewal GuARDian*), which is rapidly induced by LIF and enriched in the pluripotent epiblast. We show that *Asgard* is essential for efficient mESC self-renewal and operates through a dual mechanism: it acts as a precursor for the differentiation-promoting miRNA *Odin*, while simultaneously functioning as a sponge that sequesters *Odin* and related miRNAs. This duality positions *Asgard* as a unique integrative hub that connects STAT3 transcriptional inputs with post-transcriptional regulation of pluripotency.

## Results

### Identification of a novel LIF/STAT3 target gene as a potential regulator of pluripotency in mouse embryonic stem cells

We used E14-S3ER cells to analyze the STAT3 transcriptome in mouse ESCs (Bourillot *et al*, 2009). These cells express a hormone-inducible mutant form of STAT3 (STAT3-ER^T2^), whose activity is specifically controlled by tamoxifen (Matsuda *et al.*, 1999). E14-S3ER cells can self-renew in presence of either LIF or 4’hydroxy-tamoxifen (4’OHT). Transcriptome analysis of E14-S3ER cells deprived of 4’OHT, followed by restimulation with either LIF or 4’OHT, identified 65 genes commonly regulated by both stimuli. Among them was an uncharacterized transcript corresponding to Affymetrix probe ID 1456160_at (**Fig. 1A**). RT-PCR confirmed that *1456160_at* expression was activated by LIF or 4’OHT. A 1.5-fold upregulation upon 4’OHT stimulation in presence of cycloheximide indicated that *1456160_at* is a direct STAT3 target gene in E14-S3ER cells (**Fig. 1A**). To validate this regulation in a different context, we examined *1456160_at* expression in the E14tg2a mESC line after 24 h of LIF deprivation, followed by for 24 h of LIF restimulation. RT-PCR revealed a threefold reduction in expression after LIF withdrawal and a sevenfold induction upon LIF re-addition (**Fig. 1B**). We next assessed *1456160_at* expression during embryoid body (EB) differentiation. Its transcript levels decreased as early as day 1, similar to the downregulation of key pluripotency factors *Klf4* and *Klf5*, but were not completely extinguished during differentiation (**Fig. 1C**).

**Figure 1.**
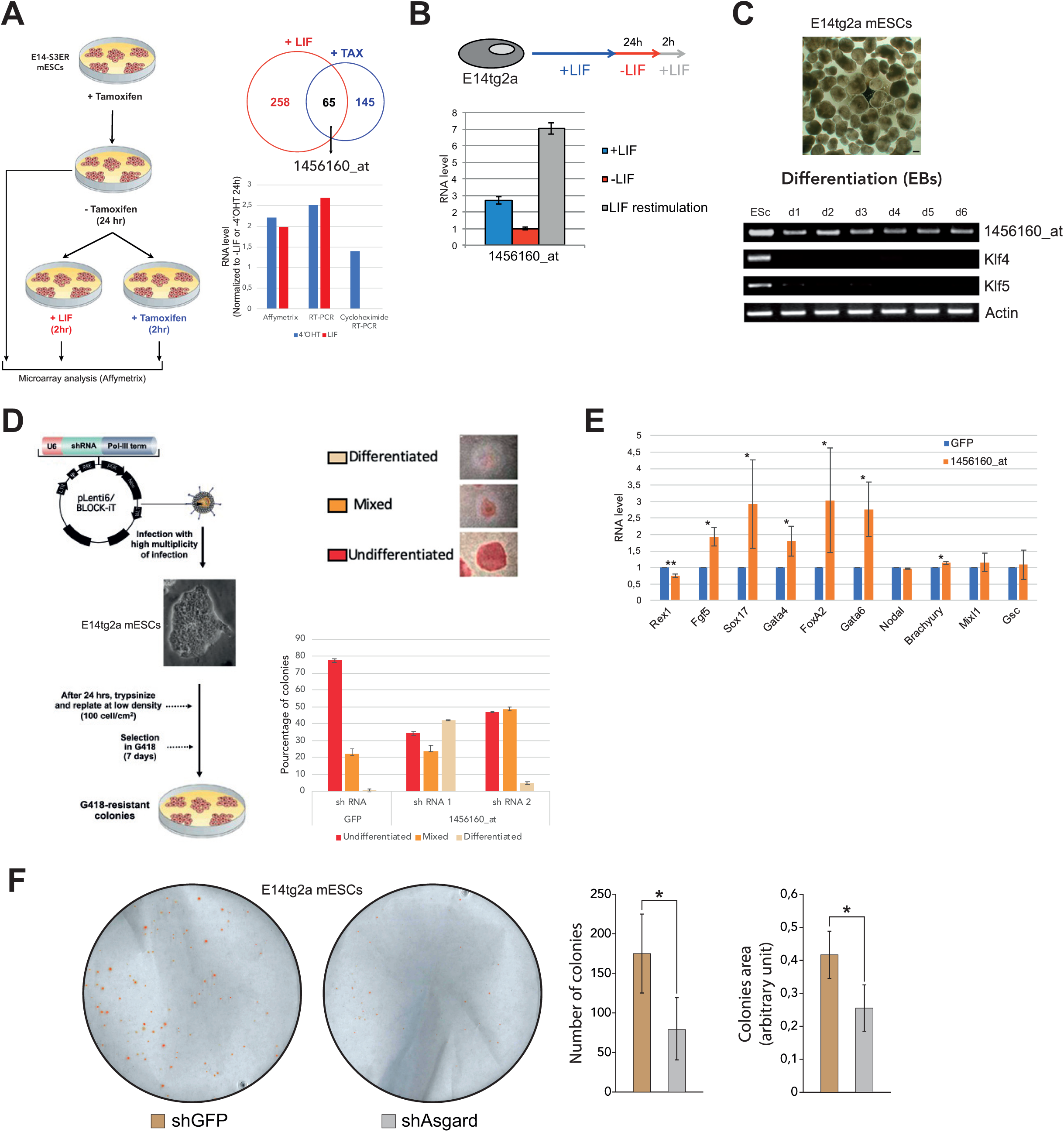
Identification of *Asgard*, a novel pluripotency regulator in mESCs. **(A)** Strategy for identifying STAT3 target genes. The Venn diagram shows genes induced by leukemia inhibitory factor (LIF), 4-hydroxytamoxifen (4’OHT), or both, including an uncharacterized gene corresponding to Affymetrix probe ID *1456160_at*. Histogram shows *1456160_at* expression in E14-S3ER cells treated with LIF or 4’OHT, measured by microarray and RT-PCR, normalized to untreated controls. **(B)** *1456160_at* mRNA levels (ΔCt) in E14tg2a cells cultured with continuous LIF, deprived of LIF for 24 h, or restimulated with LIF for 2 h, normalized to untreated control. **(C)** Phase-contrast images of embryoid bodies (EBs) at day 4 of differentiation and RT-PCR analysis of *1456160_at*, *Klf4*, *Klf5*, and *β-actin* expression during EB-mediated differentiation of E14tg2a cells. **(D)** Experimental workflow for testing *1456160_at* function in mESC pluripotency. Histogram shows the proportion of undifferentiated (red), mixed (orange), and differentiated (beige) colonies in clonal assays after knockdown with two independent shRNAs (*pLenti6/BLOCK-iT-PGKneoR*). **(E)** qRT-PCR analysis of pluripotency and lineage markers (*Rex1*, *Fgf5*, *Sox17*, *Gata4*, *FoxA2*, *Gata6*, *Nodal*, *Brachyury*, *Mixl1*, *Gsc*) in colonies from panel D. **(F)** Knockdown of *1456160_at* (*Asgard*) in E14tg2a cells using shRNAs targeting regions distinct from the Affymetrix probe sequence. Histograms show colony number and area in clonal assays after *Asgard* knockdown. Data represent means ± SE from three independent experiments; Student’s *t*-test (\**p* < 0.05; \*\**p* < 0.01; \*\*\**p* < 0.001).

To explore the functional role of *1456160_at* in ESC self-renewal, we performed knockdown experiments using lentiviral vectors encoding two independent small hairpin (sh)RNA targeting*1456160_at* (*sh1456160_at-1 and sh1456160_at-2*), or a control shRNA targeting *GFP,* together with a neomycin resistant cassette. Twenty-four hours post-infection, E14tg2a ES cells were replated at clonal density and cultured for 7 days under G418 selection to eliminate non-infected cells. Knockdown efficiency was confirmed by RT-PCR (**Sup. Fig. 1**). Colony-forming assays scored G418-resistant colonies as undifferentiated (alkaline phosphatase (AP)^+^, mixed (AP**^+^**/AP**^-^**), or differentiated (AP**^-^**). Control cells expressing *shGFP* maintained nearly 80% undifferentiated AP**^+^** colonies. In contrast, *sh1456160_at-1* and *sh1456160_at-2* reduced the proportion of AP**^+^** colonies by ∼ 45% and ∼ 30%, respectively, with a corresponding increase in mixed AP**^+^**/AP**^-^**colonies (*sh1456160_at-1,* +5%*; sh1456160_at-2*, +30%) and AP**^-^**differentiated colonies (*sh1456160_at-1*, +40%, *sh1456160_at-2*, +5%) (**Fig. 1D**). qRT-PCR analysis of pooled AP^+^, mixed, and AP**^-^** colonies revealed that *1456160_at* knockdown reduced expression of the pluripotency marker *Rex1* (*Zfp42*), while inducing early ectoderm (*Fgf5*) and endoderm markers (*Sox17*, *FoxA2*, *Gata4* and *Gata6*) (**Fig. 1E**). Collectively, these findings identify *1456160_at* as a novel LIF/STAT3 target gene that promotes pluripotency and inhibits differentiation in mESCs. Based on its role, we renamed it *Asgard* (Another Self-renewal GuARDian).

### Asgard is a long non-coding RNA regulating mESC self-renewal

The Affymetrix probe ID 1456160_at corresponds to a 407-nucleotide expressed sequence tag (EST) located on chromosome 11, in the antisense orientation within the first intron of the *Nucleoredoxin* (*Nxn*) gene (**Fig. 2A**). To determine the full-length sequence of the *Asgard* transcript, we performed RACE-PCR, obtaining a 766-nt fragment that was subsequently sequenced (**Sup. Fig. 2A**). Given its intronic location, we hypothesized that *Asgard* encodes a non-coding RNA. Northern blot analysis of RNA from wild-type E14tg2a mESCs–as well as from Dicer^-/-^ and DGCR8^-/-^ mESCs defective in small RNA biogenesis (Kanellopoulou *et al*, 2005; Wang *et al.*, 2007)–was performed using a probe derived from the 766-nt fragment. Northern blotting revealed a ∼6,000-nt transcript corresponding to *Asgard*, with low expression in wild-type cells but markedly increased levels in Dicer⁻/⁻ and DGCR8⁻/⁻ mESCs (**Fig. 2B**). This increase was further supported by transcriptome analysis of Dicer⁻/⁻ mESCs (**Fig. 2E**). These data suggest that *Asgard* is a lncRNA subject to processing by Dicer and DGCR8. The complete *Asgard* sequence was reconstructed by sequencing RT-PCR products amplified with progressively extended 3′ and 5′ primers beyond the region defined by the initial RACE-PCR fragment (**Sup. Fig. 2B**). Sequence analysis using the NCBI ORFfinder tool (https://www.ncbi.nlm.nih.gov/orffinder/) detected no long open reading frames, further supporting its classification as a lncRNA (**Sup. Fig. 4A**).

**Figure 2.**
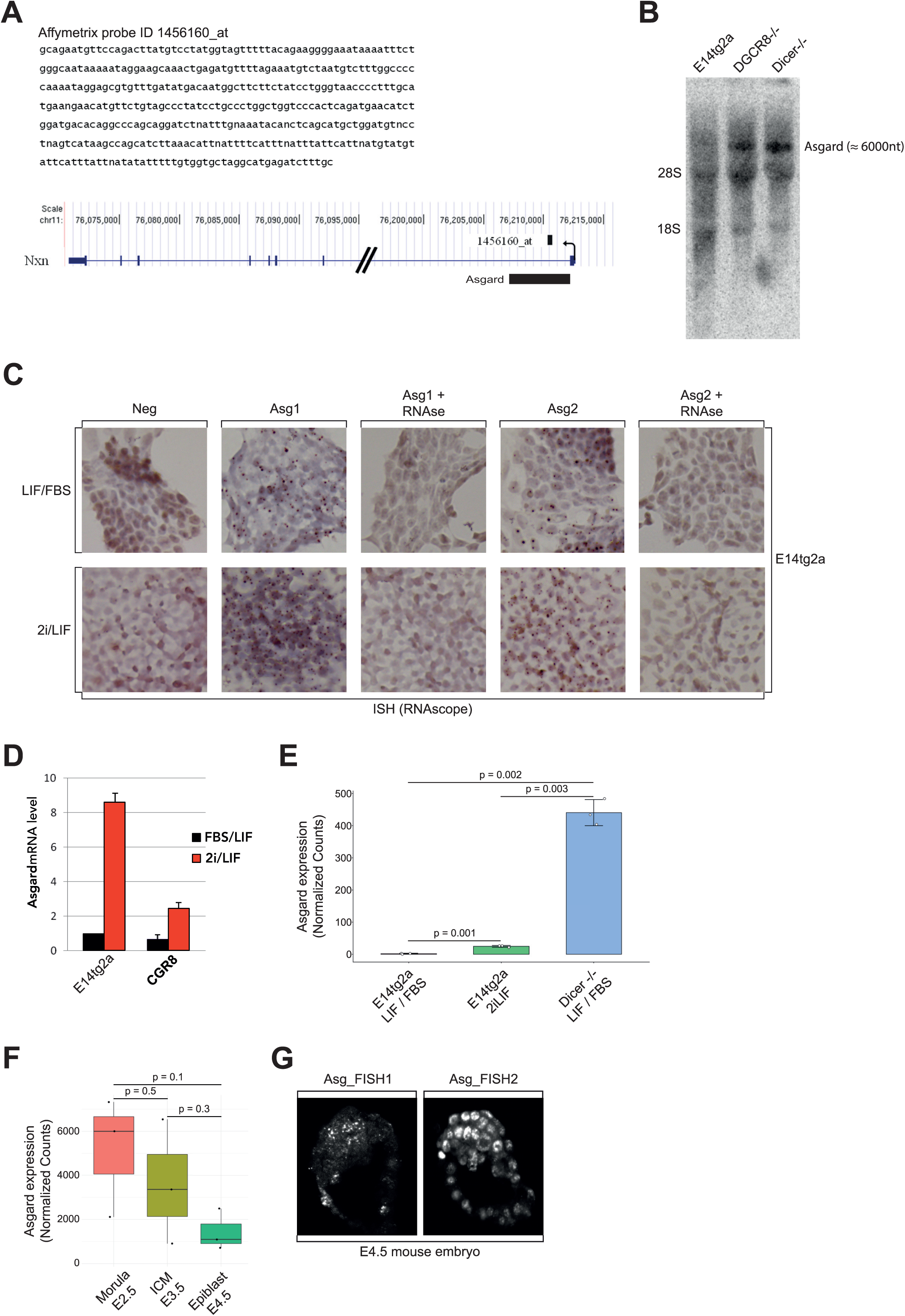
Expression pattern of *Asgard* in mESCs. **(A)** Sequence and genomic localization of Affymetrix probe ID *1456160_at* within the *Asgard* locus, visualized using the UCSC Genome Browser (https://genome-euro.ucsc.edu). **(B)** Northern blot analysis of *Asgard* expression in wild-type (E14tg2a), *DGCR8^-/-^*, and *Dicer^-/-^* mESCs. **(C)** *In situ* hybridization of E14tg2a cells cultured in LIF/serum or 2i/LIF using two independent RNAscope probes targeting *Asgard*. **(D)** qRT-PCR analysis of *Asgard* mRNA levels (ΔCt) in E14tg2a and CGR8 cells cultured in LIF/serum or 2i/LIF, normalized to LIF/serum. **(E)** Histogram representation of the mRNA level (DESeq2-normalized counts) of *Asgard*, in E14tg2a mESCs cultured in LIF/serum or 2I/LIF, and *Dicer^-/-^* mESCs. **(F)** Boxplot representation of the mRNA level (DESeq2-normalized counts) of *Asgard*, in mouse morula (E.2.5), mouse ICM (E3.5) and mouse epiblast (E4.5). **(G)** Fluorescent *in situ* hybridization (FISH) detection of *Asgard* in E4.5 mouse embryos using two distinct FISH probes. Statistical test: t-test.

We next examined *Asgard* expression under different culture conditions. RNA *in situ* hybridization using two independent probes spanning the full transcript revealed higher expression in E14tg2a mESCs cultured in 2i/LIF compared to serum/LIF conditions (**Fig. 2C**). This enrichment was confirmed by qRT-PCR, showing an ∼8-fold increase in E14tg2a cells and a ∼4-fold increase in CGR8 cells under 2i/LIF (**Fig. 2D**), as well as by analysis of published RNA-seq datasets (**Fig. 2E, Sup. Fig. 2C**). Given that mESCs correspond to the preimplantation epiblast *in vivo*, we analyzed *Asgard* expression during early embryogenesis using published RNA-seq datasets. Expression was detected at the morula (E2.5), inner cell mass (E3.5), and epiblast (E4.5) stages, with a trend toward higher levels at earlier stages, although this difference did not reach statistical significance (**Fig. 2F, Sup. Fig. 2D**). Consistent with these data, fluorescent in situ hybridization (FISH) on E4.5 embryos using two independent probes revealed enrichment of *Asgard* in the epiblast compared to surrounding tissues (**Fig. 2G**), supporting its association with the naïve pluripotent state.

To investigate *Asgard* function, we knocked down its expression using lentiviral shRNAs targeting sequences outside the original 1456160_at probe region (*pLenti6/BLOCK-iT-PGKneo-shAsgard*). Twenty-four hours post-infection, E14tg2a mESCs were replated at clonal density and cultured for 7 days under selection. Colonies were then fixed and stained for AP, and quantified. *Asgard* knockdown led to a marked reduction in both the number and size of AP-positive colonies (**Fig. 1F**), indicating a requirement for *Asgard* in maintaining mESC self-renewal.

### *Asgard* promoter is regulated by STAT3 in mouse embryonic stem cells

To investigate the promoter activity of the *Asgard* gene, we examined ENCODE datasets (https://www.encodeproject.org). In the 5’ region of the gene, we identified a 630 bp sequence bearing characteristic promoter-associated epigenetic marks: strong RNA polymerase II occupancy, pronounced H3K4me3 enrichment, modest H3K27me3 signal, and no detectable H3K9me3 (Bernstein *et al*, 2006). This sequence, hereafter referred to as the “Sense promoter”, was cloned upstream of a luciferase reporter in the *pGL4* (*pAsg-Sense-Luc*) (**Fig. 3A, Sup. Fig. 3A**). For comparison, a 3’ region displaying similar epigenetic features–termed the “Antisense promoter”–was cloned in the same configuration (*pAsg-Antisense-Luc*) (**Fig. 3A, Sup. Fig. 3A**). E14tg2a mESCs were transfected with *pGL4-Luc* (empty vector), *pAsg-Sense-Luc*, or *pAsg-Antisense-Luc* and cultured in three conditions: continuous LIF, 24 h LIF withdrawal, or LIF withdrawal followed by 2h LIF restimulation. Luciferase assays revealed that only the Sense promoter exhibited LIF-dependent activity, which decreased 1.7-fold upon LIF removal and increased 1.9-fold upon LIF restimulation (**Fig. 3B**). Similar results were obtained in E14-S3ER mESCs, which express a hormone-inducible STAT3-ER^T2^ variant that supports proliferation in the presence of 4’OHT. 4’OHT withdrawal and re-addition confirmed that the 630 bp Sense promoter mediates STAT3-dependent luciferase activation in E14-S3ER cells (**Fig. 3C**).

**Figure 3.**
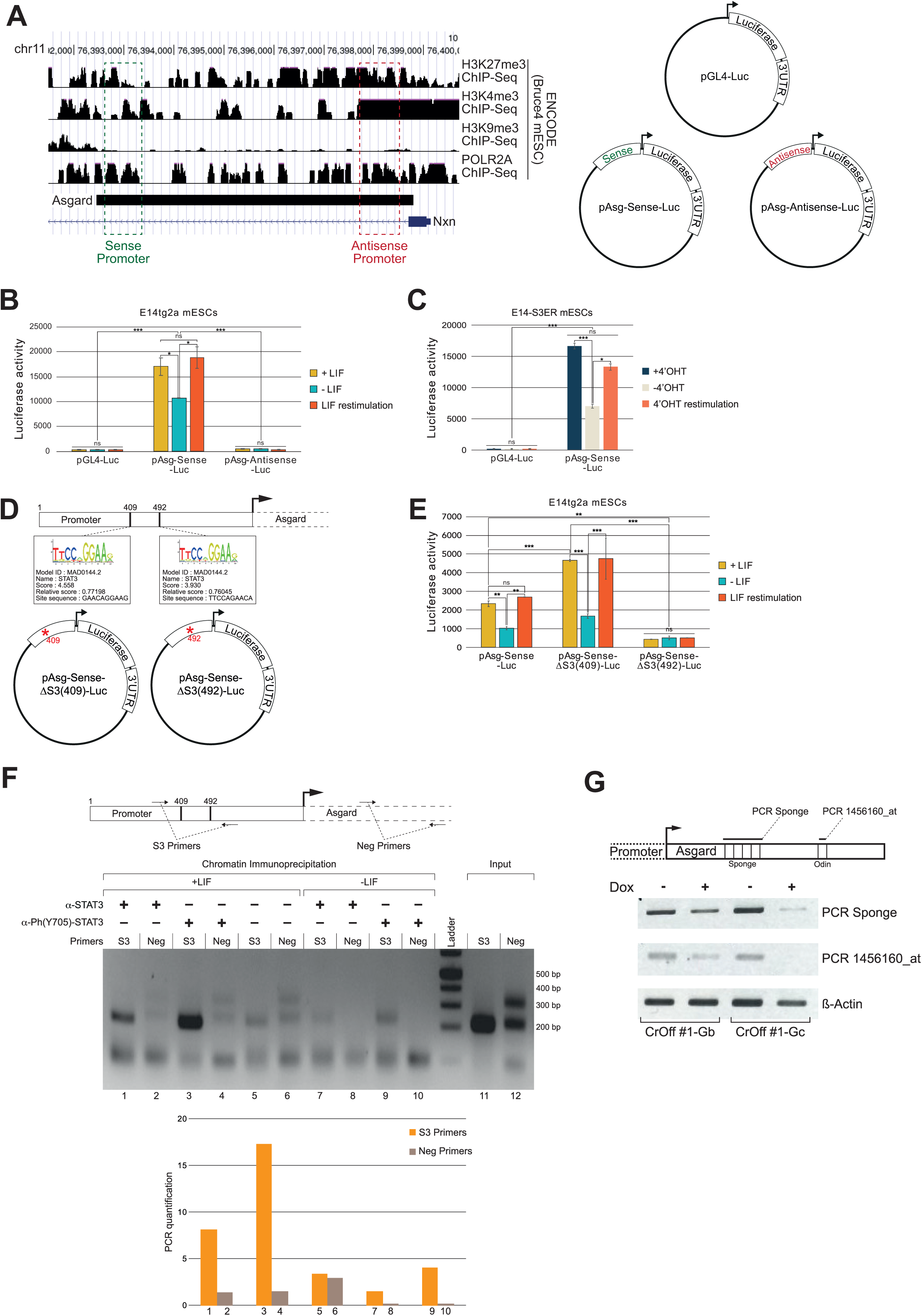
Characterisation of the *Asgard* promoter. **(A)** Chromatin landscape of the *Asgard* locus in Bruce4 mESCs. ChIP-seq binding profiles for RNA Polymerase II (Pol II), H3K4me3 (active promoter mark), and H3K9me3/H3K27me3 (repressive marks) were obtained from the ENCODE database and aligned to the *Asgard* genomic region. Two distinct sequences showing promoter-associated epigenetic features were identified: a 5′ “Sense Promoter” and a 3′ “Antisense Promoter”. Both sequences were cloned into the *pGL4-Luc* reporter plasmid. Schematic shows *pGL4-Luc, pAsg-Sense-Luc,* and *pAsg-Antisense-Luc* constructs. **(B)** Luciferase reporter assay in E14tg2a mESCs transfected with *pGL4-Luc, pAsg-Sense-Luc*, or *pAsg-Antisense-Luc*, under continuous LIF, 24 h LIF deprivation, or 2 h LIF re-stimulation. **(C)** Luciferase activity in E14-S3ER mESCs transfected with *pGL4-Luc* or *pAsg-Sense-Luc* under continuous 4’OHT, 24-h 4’OHT withdrawal, or 2 h 4’OHT re-stimulation. **(D)** *In silico* prediction of two STAT3 binding sites at positions 409 and 492 of the Sense Promoter (JASPAR database). Schematic of mutant constructs *pAsg-Sense-ΔS3(409)-Luc* and *pAsg-Sense-ΔS3(492)-Luc*, with individual STAT3 site mutation. **(E)** Luciferase assay in E14tg2a mESCs transfected with wild-type (*pAsg-Sense-Luc*) or mutant (*pAsg-Sense-ΔS3(409)-Luc* and *pAsg-Sense-ΔS3(492)-Luc*) reporter plasmids, under LIF, 24 h LIF deprivation, or 2 h LIF re-stimulation. **(F)** ChIP-PCR analysis in E14tg2a mESCs using antibodies to total STAT3 or phosphorylated STAT3 (Y705). Primers targeted the *Asgard* promoter (S3) or a negative control region (Neg). Left: representative electrophoresis gel image; Right: quantification of PCR signal. **(G)** RT-PCR analysis of *Asgard* expression in CrOff #1-Gb and CrOff #1-Gc cells under –doxycycline and +doxycycline conditions. The positions of the PCR products along the *Asgard* gene are indicated in the schematic shown above. Data represent mean ± SE from three independent experiments. \**p* < 0.05; \*\**p* < 0.01; \*\*\**p* < 0.001 (Student’s *t*-test).

JASPAR analysis (https://jaspar.elixir.no) identified two putative STAT3-binding sites within the Sense promoter at positions 409 and 492 (**Fig. 3D**). To assess their functional relevance in regards to the activation of the *Asgard* gene in response to LIF, each site was mutated by site-directed mutagenesis, generating *pAsg-Sense-ΔS3(409)-Luc* and *pAsg-Sense-ΔS3(492)-Luc*) (**Fig. 3D, Sup. Fig. 3B**). After transfection into E14tg2a cells, luciferase assays revealed that mutation of the 492 site abolished LIF-induced promoter activation, whereas mutation of the 409 site enhanced the response to LIF by ∼ 2-fold (**Fig. 3E**). Similar results were obtained in E14-S3ER cells cultured with 4’OHT (**Sup. Fig. 3C**).

To test whether STAT3 directly binds the *Asgard* promoter, we conducted ChIP-PCR in mESCs cultured with (+LIF) or without (-LIF, 24h) LIF. LIF stimulation induces phosphorylation of STAT3 on tyrosine 705, promoting its dimerization, nuclear translocation, and binding to the promoters of target genes (Zhong *et al*, 1994). Chromatin was immunoprecipitated using antibodies against total STAT3 or STAT3 phosphorylated on Tyr705 [P(Y705)-STAT3]. Primers flanking the STAT3-binding sites in *Asgard* promoter (S3 primers) and control primers targeting a STAT3-free region (neg primers) were used for PCR. In +LIF conditions, STAT3 binding was detected with both antibodies, with 2.5-fold (STAT3) and 5.5-fold [P(Y705)-STAT3] enrichment relative to no-antibody control (**Fig. 3F**, lanes 1 and 3 vs. lane 5). No enrichment was observed with neg primers (lanes 2, and 4 vs. lane 6). LIF withdrawal reduced STAT3 binding by ∼ 4-fold (lane 7 vs. lane 1; lane 9 vs. lane 3), confirming LIF-dependent recruitment of STAT3 to the *Asgard* promoter region containing binding sites 1 and 2.

To functionally validate the 630 bp Sense region as the *Asgard* promoter, we employed a CRISPR-based transcriptional repression approach (Gil *et al.,* 2024). E14tg2a mESCs were engineered to express a doxycycline-inducible dCas9 fusion protein coupled to DNMT3A, DNMT3L, and the KRAB repressive domain (**Sup. Fig. 3D**), generating the CrOff #1 cell line. These cells were subsequently transfected with guide RNAs targeting the 630 bp Sense region (**Sup. Fig. 3E**) and subjected to hygromycin selection. Following doxycycline induction, *Asgard* expression was assessed by RT-PCR. Analysis of two independent clones, CrOff #1-Gb and CrOff #1-Gc, generated using guide RNAs b and c, respectively, revealed a consistent reduction in *Asgard* expression under +Dox conditions compared to −Dox controls (**Fig. 2G**). This decrease was observed with two primer pairs tested, which specifically amplify either the *Odin* miRNA or the sponge domain, and was more pronounced in the CrOff #1-Gc clone. Notably, both the sponge domain and *Odin* miRNA levels were concomitantly reduced. These results demonstrate that the 630 bp Sense region functions as the *Asgard* promoter and drives the coordinated expression of both the *Odin* miRNA and the sponge domain, indicating that they originate from a common transcriptional unit.

Together, these results demonstrate that *Asgard* expression is regulated by LIF-dependent STAT3 binding to its Sense promoter, with the site at position 492 being essential for activation.

### Long non-coding RNA *Asgard* functions both as a microRNA precursor and a microRNA sponge

The regulation of *Asgard* by Dicer and DGCR8 suggested that it may serve as a primary microRNA (pri-miRNA). Sequence analysis using the miRortho database (http://cegg.unige.ch/mirortho/) revealed a region with strong homology to microRNA *miR-1195*, with a highly significant e-value of 5.7 × 10⁻¹⁴ (**Fig. 4A**). Further analysis with the MirID algorithm (http://datalab.njit.edu/RNAcenter/) classified this region as a putative pre-miRNA (**Sup. Fig. 4B**). Using the stem-loop RT-PCR protocol developed by Varkonyi-Gasic et al., we successfully detected the expression of the miRNA encoded by *Asgard* (**Fig. 4B**) (Varkonyi-Gasic *et al*, 2007). To assess its function, synthetic miRNA mimics were transfected into E14tg2a mESCs. qRT-PCR analysis showed downregulation of the pluripotency markers *Nanog* and *Rex1*, along with upregulation of the early ectodermal marker *Fgf5* and various mesodermal and endodermal markers (**Fig. 4C**). Overexpression of the miRNA via lentiviral transduction (OE-miRAsgard) and subsequent neomycin selection for 7 days yielded cells with a differentiated morphology. qRT-PCR confirmed reduced pluripotency marker expression and increased differentiation marker expression (**Fig. 4D**). These data indicate that the *Asgard*-encoded miRNA promotes differentiation. This miRNA was designated *Odin* (*One Differentiation INducer*).

**Figure 4.**
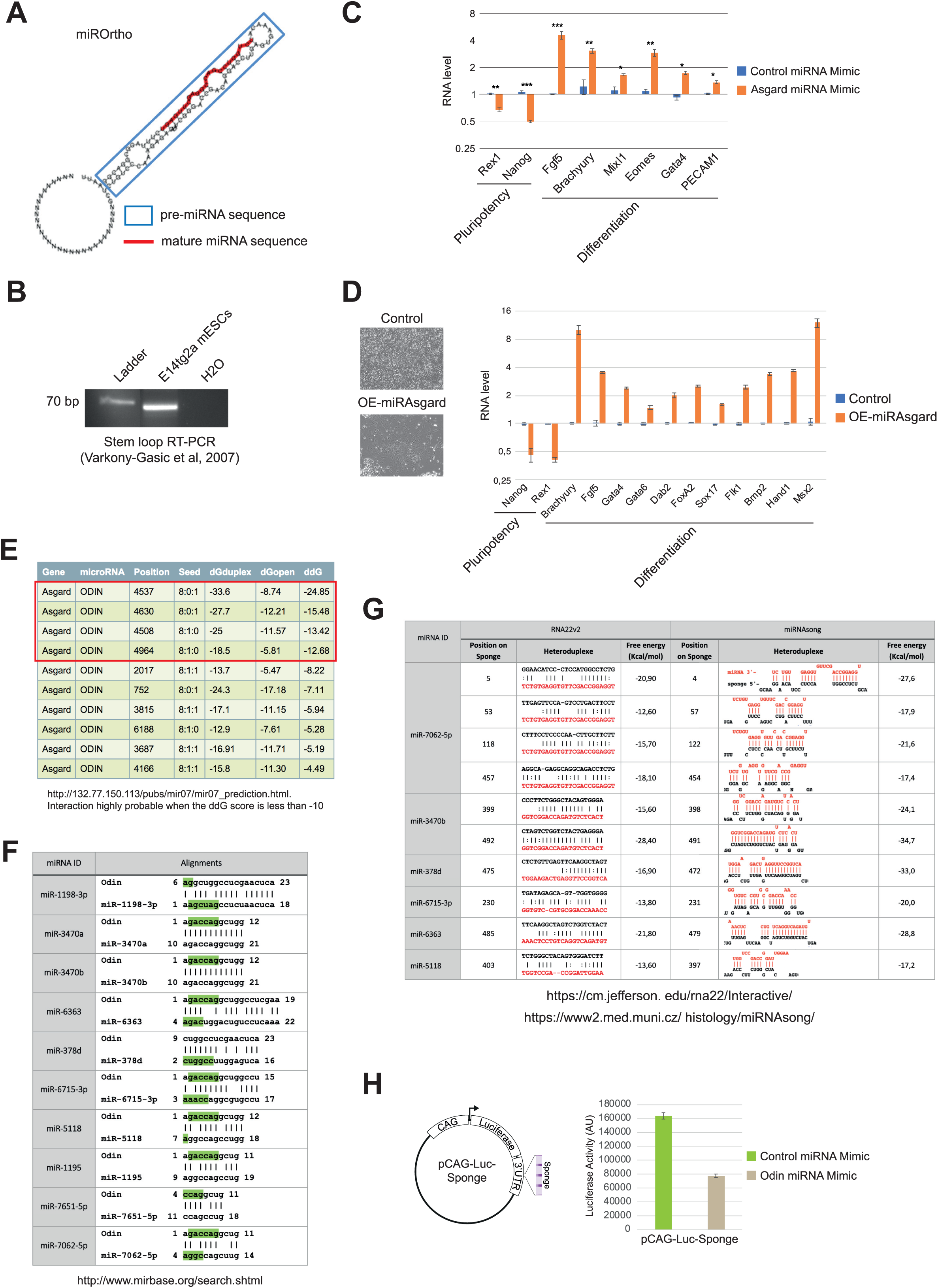
The lncRNA *Asgard* functions as a miRNA sponge. **(A)** *In silico* prediction of a pre-miRNA hairpin and mature miRNA sequence within the *Asgard* gene (miROrtho database). **(B)** Detection of mature *Asgard* miRNA in E14tg2a mESCs cultured in LIF/FBS using stem-loop RT-PCR. **(C)** qRT-PCR analysis of pluripotency (*Rex1*, *Nanog*) and differentiation (*Fgf5*, *Brachyury*, *Mixl1*, *Eomes*, *Gata6*, and *PECAM1*) markers in E14tg2a mESCs transfected with an *Asgard* miRNA mimic or control mimic. **(D)** Phase-contrast microscopy of E14tg2a mESCs overexpressing *Asgard* miRNA (OE-miRAsg) versus control cells. qRT-PCR analysis of pluripotency (*Nanog*, *Rex1*) and differentiation (*Brachyury*, *Fgf5*, *Gata4*, *Gata6*, *Dab2*, *FoxA2*, *Sox17*, *Flk1*, *Bmp2*, *Hand1*, *Msx2*) markers in OE-miRAsg and control cells. **(E)** *In silico* prediction of miRNA response elements (MREs) for *Odin* within *Asgard* (Seagal database). **(F)** Identification of miRNAs homologous to the predicted *Odin* miRNA by aligning the *Odin* sequence against the mouse miRNA database (miRBase BLAST). Seed sequences are underlined in green. **(G)** Predicted binding of candidate miRNAs (*miR-7062-5p, miR-3470b, miR-378d, miR-6715-3p, miR-6363*, and *miR-5118*) to the *Asgard* sponge domain using RNA22v2 and miRNAsong. Free binding energies (kcal/mol) are indicated for each interaction. **(H)** Schematic of luciferase reporter construct *pCAG-Luc-Sponge*, in which the *Asgard* sponge domain was cloned into the 3′ UTR of luciferase under the CAG promoter. Luciferase activity was measured in E14tg2a mESCs co-transfected with *pCAG-Luc-Sponge* and either an *Asgard* miRNA mimic or a control mimic.

The function of *Odin* as a differentiation inducer contrasts with the function of *Asgard* as a pluripotency stabilizer. To address this issue, we analyzed potential *Odin* targets using Segal lab tools (http://132.77.150.113/pubs/mir07/mir07_prediction.html), which revealed four putative Odin-binding sites clustered within a 500-nt region of the *Asgard* sequence (**Fig. 4E**). This suggested the existence of a “sponge domain” capable of sequestering *Odin* or related miRNAs. BLASTn searches against all annotated mouse miRNAs in miRBase release 22 (Kozomara *et al*, 2019) identified 10 miRNAs with sequence similarity to *Odin*: *miR-1195*, *miR-5118, miR-1198-3p, miR-3470a, miR-3470b, miR-7062-5p, miR-6363, miR-7651-5p, miR-378d*, and *miR-6715-3p* (**Fig. 4E**). To predict binding sites for these *Odin*-related miRNAs within the *Asgard* sponge domain, we used two independent prediction tools–RNA22v2 (https://cm.jefferson.edu/rna22/Interactive/) (Miranda *et al*, 2006) and miRNAsong (https://www2.med.muni.cz/histology/miRNAsong/) (Barta *et al*, 2016)–and retained only sites predicted by both. Six miRNAs were predicted to bind the sponge domain: *miR-7062-5p* (four predicted binding sites), *miR-3470b* (two sites), and one site each for *miR-378d, miR-6715-3p, miR-6363*, and *miR-5118* (**Fig. 4G**). These predictions support the hypothesis that *Asgard* sequesters a subset of *Odin*-related miRNAs, particularly *miR-7062-5p* and *miR-3470b*. To functionally test this, the 500-nt sponge domain was cloned into the 3′ untranslated region (3′UTR) of the luciferase gene in *pCAG-Luc*, generating *pCAG-Luc-Sponge*. This construct was co-transfected into E14tg2a mESCs with either a negative control mimic or a synthetic *Odin* mimic. Luciferase assays revealed that the *Odin* mimic reduced reporter activity by 2-fold compared to the control (**Fig. 4H**), providing functional evidence that *Asgard* acts as a miRNA sponge, capable of sequestering *Odin* and related miRNAs to modulate their regulatory effects.

### Asgard sponge domain sustains mESC self-renewal

We used the miRWalk interface (http://mirwalk.umm.uni-heidelberg.de/search_mirnas/)(Sticht *et al*, 2018) to identify predicted target genes of *miR-7062-5p, miR-3470b, miR-378d, miR-6715-3p, miR-6363*, and *miR-5118*, followed by KEGG pathway enrichment analysis. *Odin,* together with these miRNAs, was predicted to target genes enriched in pathways essential for mESC pluripotency, including PI3K-AKT, JAK-STAT, MAPK, Wnt, p53, and canonical pluripotency networks (**Fig. 5A, Sup. Fig. 5A**). As most of these miRNAs are predicted to promote mESC differentiation, we hypothesized that *Asgard* may preserve self-renewal by sequestering them. To test this, we overexpressed the *500-nt* sponge domain in mESCs using the *pPB_Sponge* plasmid. Two independent clones (E14^Sponge#1^ and E14^Sponge#2^) showed marked resistance to differentiation in LIF dilution and deprivation assays (**Fig. 5B, Sup. Fig. 5B**). Whereas control cells formed AP^+^ colonies only at high LIF concentrations (> 100 U/mL), both E14^Sponge#1^ and E14^Sponge#2^ maintained AP^+^ colonies at concentrations as low as 0.1 U/mL. Moreover, they retained AP^+^ colony formation after 48 h of LIF-deprivation, while control cells lost this capacity after just 24 h. Furthermore, colony counts—a measure of clonogenic self-renewal—were significantly higher in sponge-overexpressing clones than in controls. At 1000 U/mL LIF, E14^Sponge#1^ and E14^Sponge#2^ generated 118 ± 5 and 104 ± 12 colonies, respectively, compared to 23 ± 6 in control E14tg2a cells (**Fig. 5C**). E14^Sponge#1^ consistently exhibited the greatest clonogenic capacity, both in the presence and absence of LIF, and this enhancement persisted after prolonged LIF deprivation (**Fig. 5D**).

**Figure 5.**
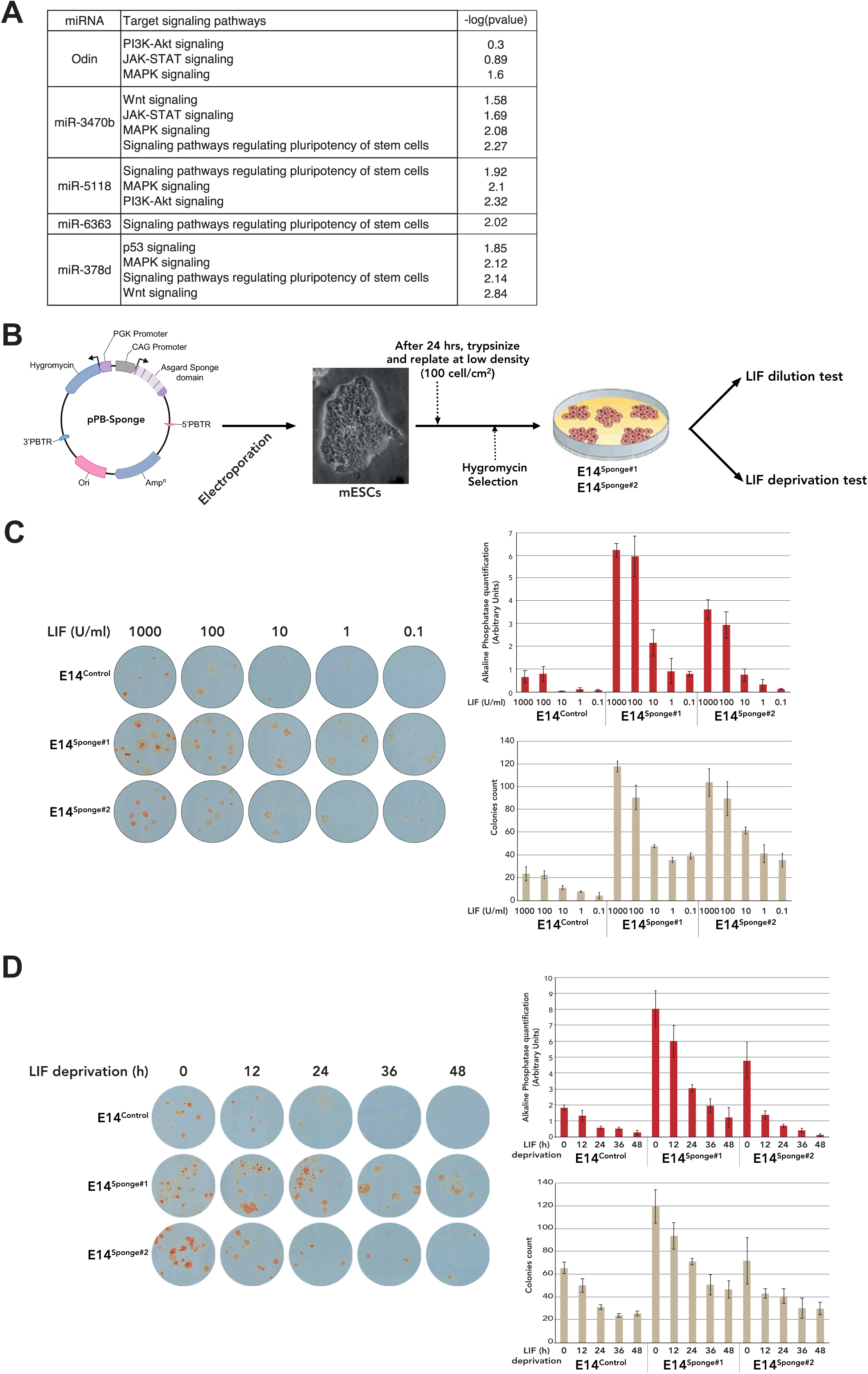
Overexpression of the *Asgard* sponge domain promotes mESC self-renewal under suboptimal culture conditions. **(A)** Predicted target signaling pathways of *Odin, miR-3470b, miR-5118, miR-6363*, and *miR-378d*–miRNAs potentially regulated by the *Asgard* sponge domain and known to modulate mESC pluripotency. **(B)** Schematic of the experimental strategy for functional analysis of the *Asgard* sponge domain. The sponge sequence was cloned into a *pPB-Hygromycin* PiggyBac vector under the constitutive CAG promoter (*pPB-Sponge*) and stably integrated into E14tg2a mESCs. **(C)** Alkaline phosphatase (AP) staining of E14^Control^ cells and two independent *Asgard* sponge-expressing clones (E14^Sponge#1^ and E14^Sponge#2^) cultured at clonal density in media containing decreasing LIF concentrations. Right: quantification of AP^+^ colonies and total colony number. **(D)** AP staining of E14^Control^, E14^Sponge#1^, and E14^Sponge#2^ cells cultured at clonal density without LIF for 0, 12, 24, 36, and 48 h, followed by re-stimulation with 1000 U/mL LIF for one week. Right: quantification of AP^+^ colonies and total colony number in the clonal assay.

Collectively, these findings indicate that the *Asgard* sponge domain enhances the robustness of pluripotency by buffering differentiation-promoting miRNAs. We propose a model in which a novel LIF target gene encodes a lncRNA that both generates and sequesters specific miRNAs, thereby fine-tuning the balance between self-renewal and differentiation (**Fig. 6**).

**Figure 6.**
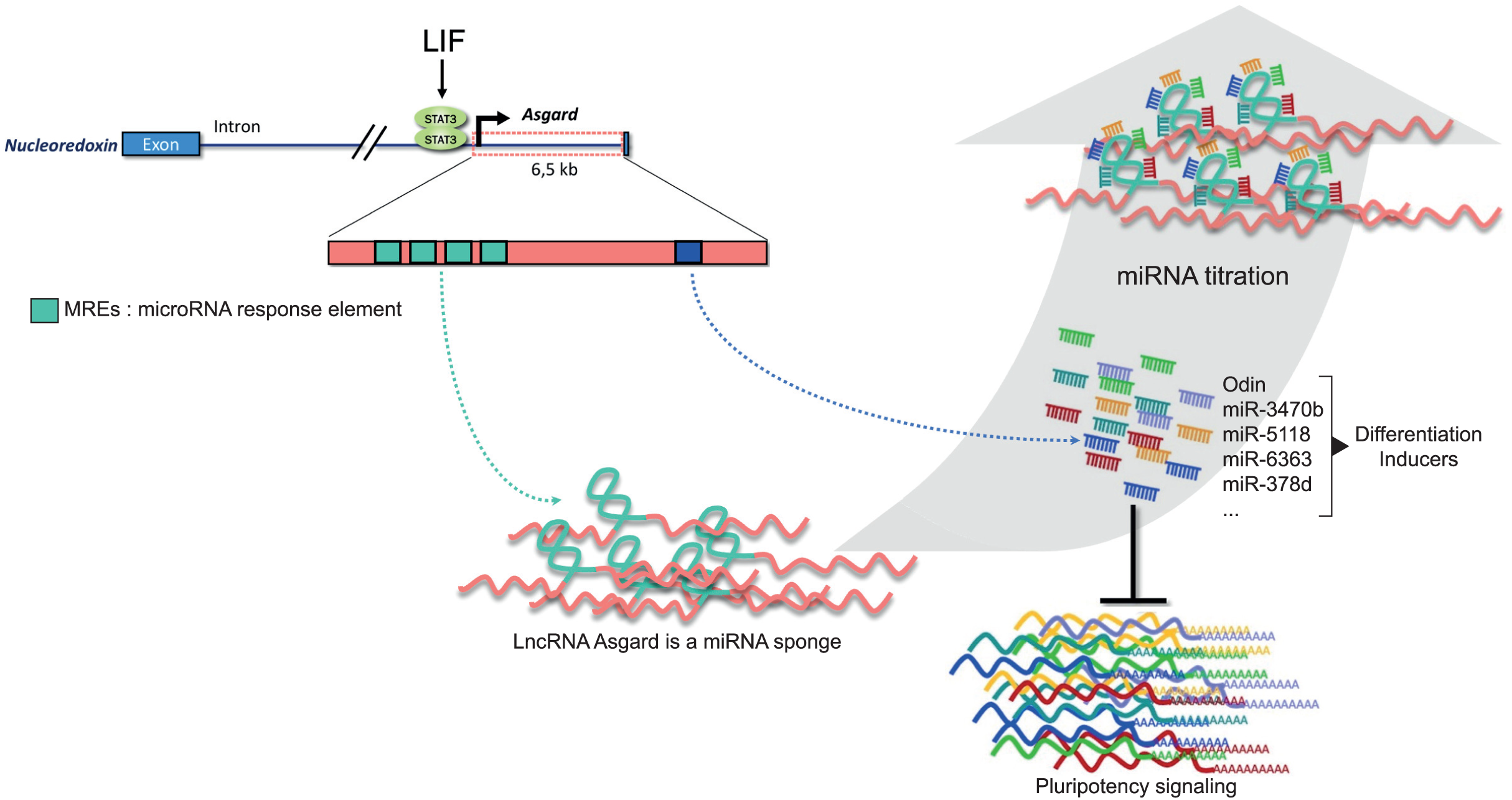
Proposed model for the role of the lncRNA *Asgard* in supporting mESC pluripotency. Schematic representation of the regulatory function of *Asgard* in maintaining pluripotency in mouse embryonic stem cell (mESCs). *Asgard* is transcriptionally activated by STAT3 signaling and contributes to pluripotency maintenance by acting as a molecular sponge for multiple microRNAs, including *Odin* and *miR-3470b*, which are predicted to target components of key pluripotency-associated signaling pathways. Through this dual mechanism—STAT3 dependent activation and miRNA sequestration—*Asgard* helps stabilize the pluripotent state.

## Discussion

In this study, we identified *Asgard*, a previously uncharacterized lncRNA, as a direct transcriptional target of STAT3 and a novel regulator of pluripotency in mESCs. *Asgard* is rapidly induced upon LIF stimulation, and its depletion compromises self-renewal while promoting differentiation. Strikingly, *Asgard* integrates two antagonistic functions: it acts as a precursor for the differentiation-inducing microRNA *Odin*, while simultaneously functioning as a sponge that titrates *Odin* and related miRNAs away from their targets. This dual activity underscores a unique principle in stem cell regulation, where a single lncRNA transcript can both generate differentiation pressure and buffer it, thereby stabilizing the pluripotent state.

Our findings extend the canonical view of STAT3 signaling in pluripotency. While STAT3 is well established as a master regulator of the transcriptional network that sustains self-renewal (Matsuda *et al.*, 1999; Niwa *et al.*, 2009), its role in controlling post-transcriptional layers of regulation has remained less clear. The discovery of *Asgard* highlights that STAT3 also reinforces pluripotency through non-coding RNA circuits, in which lncRNAs and miRNAs form intertwined feedback loops. Similar principles have been described for *Lncenc1*, which sustains pluripotency by sponging miR-128 to stabilize *Klf4* expression (Monteleone *et al.*, 2024). However, *Asgard* differs by embodying both miRNA precursor and lncRNA functions within the same transcript, positioning it as a self-contained regulatory module. The integration of miRNA and lncRNA functions within *Asgard* may provide an evolutionary advantage by ensuring robustness of pluripotency networks. On one side, the processing of *Asgard* into *Odin* miRNA provides a latent differentiation cue, priming the system for rapid lineage commitment when external signals reduce STAT3 activity. On the other, the sponge activity of *Asgard* buffers pluripotency factors from premature repression by *Odin* and related miRNAs. This architecture creates a fail-safe mechanism, in which pluripotency is maintained unless differentiation signals persistently outweigh the buffering capacity of the lncRNA. Such duality illustrates how lncRNAs can operate not merely as passive sponges or precursors, but as integrative hubs that coordinate multiple layers of RNA regulation.

This dual mechanism also highlights broader implications for stem cell biology. miRNA–lncRNA crosstalk has been reported in diverse contexts (Statello *et al.*, 2021), but *Asgard* provides one of the first clear examples in ESCs where both functions converge in a single transcript. It raises the intriguing possibility that other STAT3-regulated lncRNAs may employ similar strategies, coupling miRNA production with miRNA sequestration to fine-tune pluripotency. Moreover, this mechanism could be conserved in other developmental systems, or even co-opted in pathological settings such as cancer, where aberrant STAT3 signaling and ncRNA regulation are common.

Several questions remain open. The spectrum of miRNAs sequestered by *Asgard* requires further biochemical validation, as does the stoichiometric balance between its precursor and sponge functions. Whether these functions are temporally segregated—e.g., precursor activity during lineage priming and sponge activity during maintenance—also warrants investigation. *In vivo*, *Asgard’s* enrichment in the epiblast suggests a developmental role, but definitive proof will require genetic loss-of-function studies in embryos. Finally, it will be important to determine whether human pluripotent stem cells possess *Asgard*-like lncRNAs with dual regulatory activities.

In conclusion, our study identifies *Asgard* as a dual-function lncRNA that integrates miRNA biogenesis and miRNA sequestration downstream of STAT3 signaling. By generating and simultaneously buffering differentiation-promoting miRNAs, *Asgard* provides a unique molecular strategy to stabilize the naïve pluripotent state while preserving developmental responsiveness. These findings reveal a new layer of complexity in non-coding RNA regulation and highlight how lncRNAs can act as integrative nodes that connect transcriptional and post-transcriptional control of cell fate.

## Materials and Methods

### Mouse ES cell culture and embryoid body formation

Mouse embryonic stem cell (mESC) lines (E14, CGR8, *Dicer^-/-^*, and *DGCR8^-/-^*) were cultured at 37 °C in 7.5% CO₂ on 0.1% gelatin-coated plates. Cells were maintained in Glasgow’s Modified Eagle’s Medium (GMEM; Gibco, 21710025) supplemented with 10% fetal bovine serum (Invitrogen), 1% penicillin–streptomycin (Gibco, 10378016), 2 mM glutamine, 1% non-essential amino acids (Gibco, 11140035), 1 mM sodium pyruvate (Gibco, 11360), 0.1 mM β-mercaptoethanol, and 1000 U/ml LIF. Cultures were passaged every 2 days by trypsinization and replated at a 1:5–1:10 ratio, with daily medium replacement. For serum-free culture, cells were maintained in N2B27 medium (StemCells, SCS-SF-NB-02) supplemented with 1 μM PD0325901 (Stemgent, 04-0004), 3 μM CHIR99021 (Stemgent, 04-0006), and 1000 U/ml LIF. Cells were seeded at high density (1.25 × 10⁶ cells per 9.5 cm² well) and passaged using Accutase (Life Technologies, A1110501).

To induce differentiation, mESCs were cultured in suspension as hanging drops of LIF-free medium (∼100 cells per drop). After 3 days, embryoid bodies (EBs) were transferred to non-adherent Petri dishes on a rotary shaker and cultured for 7–10 days, with daily collection. RNA extraction and qPCR were performed as described below.

For LIF titration assays, mESCs were plated at clonal density and cultured for 7 days in media containing decreasing concentrations of LIF (1000, 100, 10, 1, or 0.1 U/ml). For LIF withdrawal experiments, cells were seeded at clonal density and maintained without LIF for different time intervals (17, 24, 41, or 48 h), after which cultures were re-stimulated with 1000 U/ml LIF for 7 days. At the end of both assays, colonies were fixed, stained for alkaline phosphatase (AP) activity, and quantified.

### Plasmid construction

shRNAs targeting *Asgard* were designed using the BLOCK-iT™ RNAi Designer (Thermo Fisher Scientific) and cloned into the *pENTR™/U6* vector, followed by subcloning into *pLenti6/BLOCK-iT-PGKpuroR*. The resulting *pLenti6-shRNA-puro* plasmids were used to generate lentiviral particles for mESC transduction. Knockdown efficiency was assessed by quantitative real-time PCR, and the shRNA sequences with their silencing efficiencies are shown in **Sup. Fig. 1**.

The “Sense promoter” and “Antisense promoter” sequences (**Sup. Fig. 3A**) were PCR-amplified from E14tg2a mESC genomic DNA using primers containing *XhoI* restriction sites and inserted into the *XhoI* site of *pGL4-Luc*, generating *pAsg-Sense-Luc* and *pAsg-Antisense-Luc*. Mutations in the two STAT3 binding sites were introduced into *pAsg-Sense-Luc* using the *QuikChange Site-Directed Mutagenesis Kit* (Agilent, Cat. No. 200519), following the manufacturer’s instructions. The resulting mutant constructs were designated *pAsg-Sense-ΔS3(409)-Luc* and *pAsg-Sense-ΔS3(492)-Luc*.

To generate *pOE-miR-Asgard*, the *Odin* sequence was cloned into *pENTR™/U6*, followed by subcloning into *pLenti6/BLOCK-iT-PGKpuroR*.

For *pCAG-Luc*, a CAG promoter fragment was PCR-amplified from *pPB-HygroR* using primers containing *SacI* and *XhoI* restriction sites, and subcloned into the corresponding sites of *pGL4-Luc*. The *Asgard* sponge domain sequence was PCR-amplified from E14tg2a mESC genomic DNA using primers with an *XbaI* site and inserted into *pCAG-Luc*, generating *pCAG-Luc-Sponge*.

The *Asgard* sponge domain was also amplified from E14tg2a mESC genomic DNA with primers containing *NheI* and *HpaI* restriction sites and subcloned into the corresponding sites of *pPB-HygroR*, yielding *pPB-Sponge*.

For CRISPRoff targeting, five independent sgRNA spacer sequences were designed to target the “Sense promoter” sequence using the CRISPick web portal (https://crispor.gi.ucsc.edu). Annealed sense and antisense oligonucleotides containing the appropriate BbsI overhangs were ligated into the BbsI site of the *pSB-BbsI-sgRNA* plasmid. Details of the spacer sequences are shown in **Sup. Fig. 3E**.

### RACE-PCR

For 3′ RACE, 5 μg of total RNA was reverse-transcribed at 45 °C for 1 h with AMV reverse transcriptase using the GeneRacer® Oligo dT primer (Invitrogen). Two microliters of cDNA were amplified with the kit’s 3′ nested primer and a forward primer specific for EST 1456160_at using Platinum® Taq High Fidelity (Invitrogen). PCR conditions were: 94 °C for 2 min; 5 cycles of 94 °C for 30 s, 72 °C for 2 min; 5 cycles of 94 °C for 30 s, 70 °C for 2 min; 25 cycles of 94 °C for 30 s, 65 °C for 30 s, 68 °C for 2 min; final extension 68 °C for 10 min. Products were analyzed on 2% agarose gels. For 5′ RACE, 3 μg of RNA were dephosphorylated with CIP, decapped with TAP, and ligated to the GeneRacer™ RNA oligonucleotide. Reverse transcription was performed with AMV RT and random primers. PCR was carried out with the GeneRacer™ 5′ primer and an EST 1456160_at antisense primer using Platinum® Taq High Fidelity under touchdown conditions: 94 °C for 2 min; 5 cycles of 94 °C for 30 s, 72 °C for 4 min; 5 cycles of 94 °C for 30 s, 70 °C for 4 min; 25 cycles of 94 °C for 30 s, 65 °C for 30 s, 68 °C for 4 min; final extension 68 °C for 10 min. PCR products were cloned into pCR®4-TOPO® (Invitrogen), transformed into *E. coli* Top10, and selected on LB agar containing ampicillin. Colonies were expanded, plasmids purified with QIAprep Spin Miniprep (Qiagen), and verified by restriction digestion and sequencing.

### Lentiviral production and ES cell infection

Lentiviral vectors expressing shRNAs or the Asgard microRNA were generated using the BLOCK-iT Lentiviral RNAi Expression System (Invitrogen). A total of 0.5 μg pLenti6/BLOCK-iT-PGKneor plasmid containing either shRNA or Asgard sequences was co-transfected with 1.5 μg ViraPower Packaging Mix (Invitrogen, K4975) and 6 μl Lipofectamine (Invitrogen, 11668) into 1 × 10⁶ 293T cells seeded in 2 ml medium. Viral supernatants were collected 48 h after transfection and clarified by centrifugation (3000 rpm, 10 min).

E14 mESCs (1 × 10⁴ cells) were seeded in 24-well plates and infected with viral supernatants supplemented with 6 μg/ml polybrene and LIF. After 24 h, medium was replaced with fresh ES medium. At 48 h post-infection, cells were trypsinized, replated at 1 × 10³ cells per 10-cm dish, and cultured for 7 days in ES medium supplemented with 250 μg/ml G418 (Sigma, A1720). Resistant colonies were fixed with methanol and stained with Fast Red TR salt (Sigma) and naphthol solution in Tris-buffer (0.1 M, pH 9.2) to detect alkaline phosphatase activity.

### RNA extraction, RT-PCR and qPCR

Total RNA was isolated using the RNeasy Mini Kit (Qiagen, 74106). Reverse transcription was performed with MuMLV-RT (Promega, M1701). For Asgard cloning, PCR was carried out using Phusion® High-Fidelity DNA Polymerase (NEB, M0530). qPCR was performed on cDNA using the LightCycler FastStart DNA Master SYBR Green kit (Roche, 12239264001), with expression levels normalized to β-actin.

### Transient transfection, luciferase assay and electroporation

Transient plasmid transfections were performed using Lipofectamine® 2000 (Life Technologies, 11668019) in Opti-MEM (Life Technologies, 31985088), according to the manufacturer’s instructions. Cells were collected 24–48 h post-transfection for qPCR or luciferase assays. Firefly and Renilla luciferase activities were measured using the Dual-Luciferase® Reporter Assay System (Promega, E1910) on a GloMax® Multi Detection System (Promega), and firefly activity was normalized to Renilla to control for transfection efficiency. For electroporation experiments, 1.2 × 10⁶ E14tg2a mouse ES cells were co-electroporated with 3 μg of the pPB-Sponge-HygroR vector and 3 μg of a plasmid encoding the transposase, using the Neon® Transfection System 100 μl kit (Thermo Scientific, MPK10096). Electroporation was performed with two pulses of 20 ms at 1300 V. Cells were subsequently plated at low density in 100-mm dishes and cultured for one week in medium supplemented with 100 μg/ml hygromycin B to select for stable integrations.

Sleeping Beauty (SB) transposon cell lines were generated by co-electroporating 1 × 10⁶ cells using the Neon Transfection System (Invitrogen; 1050 V, 20 ms, 2 pulses) with 2.5 μg of the SB transposon plasmid and 2.5 μg of the SB100X transposase-expressing vector *pCMV(CAT)T7-SB100* (Addgene #127909). Stable transfectants were selected for 7 days using hygromycin (200 μg/ml) for the *pSB-TRE-CRISPRoff-EF1A-TetOn* construct (Addgene #203355) and puromycin (1 μg/ml) for the *pSB-BbsI-sgRNA* vector (Addgene #203359). Individual clones were isolated and screened for CRISPRoff dCas9 expression by western blot. Positive clones were maintained under hygromycin selection (200 μg/ml). CrOff #1-sgRNA clones were further validated by RT-PCR and subsequently cultured under dual selection with hygromycin (200 μg/ml) and puromycin (1 μg/ml).

### Northern blot analysis

Total RNA (20 μg) was denatured at 65 °C and separated on 1.2% agarose gels containing 0.66% formaldehyde. RNAs were transferred overnight to Hybond-N+ membranes (Amersham) in 10× SSC, UV crosslinked, and hybridized at 68 °C with ^32P-labeled Asgard-specific probes. Membranes were washed sequentially with 1× SSC/0.1% SDS at room temperature and 0.5× SSC/0.1% SDS at 68 °C. Hybridization signals were detected by autoradiography.

### RNA *In Situ* Hybridization (RNAscope, Chromogenic Detection)

Cells were cultured on poly-D-lysine coated coverslips and fixed in 4% paraformaldehyde for 30 min at room temperature. After PBS washes, cells were dehydrated in an ethanol series (50–100%) and stored at –20°C. Coverslips were rehydrated and permeabilized with RNAscope Protease III (ACD) for 15 min at room temperature. Target-specific RNAscope probes were hybridized using the RNAscope 2.5 HD Detection Kit (ACD) following the manufacturer’s instructions. Signal amplification was performed using the recommended amplification reagents, followed by chromogenic development with DAB substrate. Nuclei were counterstained with hematoxylin for 2 min, rinsed, and coverslips mounted with aqueous mounting medium. Images were acquired using bright-field microscopy and analyzed with ImageJ/Fiji.

### Fluorescent *In Situ* Hybridization (FISH)

After flushing or dissection, embryos were fixed with 4% PFA in PBS overnight at 4°C and stored in methanol at −20°C. Embryos were rehydrated with 0.1% Tween 20 in PBS (PBT) and permeabilized with RIPA solution (150 mM NaCl, 1% Nonidet P-40, 0.5% sodium deoxycolate, 0.1% SDS, 1 mM EDTA, 50 mM Tris [pH 8.0]) for 10–20 min at room temperature (RT). Embryos were postfixed with 4% PFA + 0.2% glutaraldehyde for 10 min at RT, and prehybridized in hybridization buffer (50% formamide, 5× SSC, 1% SDS, 100 mg/ml yeast tRNA, 50 mg/ml heparin) for 1 hr at 65°C. Hybridization with probes was done overnight at 65°C. Embryos were washed twice for 30 min with 50% formamide, 5× SSC, 1% SDS, once for 20 min with 50% formamide, 2× SSC, 1% SDS for 20 min at 65°C, and then transferred to PBT. Embryos were blocked with manufacturer’s blocking solution for 1–2 hr at RT. Embryos were treated with anti-DIG or -FITC antibody conjugated to peroxidase (1/200; Roche) overnight at 4°C. Fluorescent staining with TSA-Cy3, -Cy5 kits was carried out according to the manufacturer’s instructions (Perkin Elmer).

### Bioinformatics analysis

All analyses were performed using R (version 4.5.3). Raw sequencing reads (FASTQ files) were pseudo-aligned to a modified mouse reference genome (GRCm38) supplemented with the *Asgard* sequence using the Salmon aligner (https://github.com/COMBINE-lab/salmon), run in quasi-mapping mode with sequence-specific bias correction and GC bias correction enabled. Transcript quantification was performed at default settings. Gene-level quantification and normalization were carried out with the DESeq2 package (version 1.48.2) (Love et al., 2014). Publicly available datasets used in this study include GSE215141, GSE172282, GSE263060, and E-MTAB-2958.

## Acknowledgements

This work was supported by the Fondation ARC pour la Recherche sur le Cancer (PJA-20151203436 to P-Y.B.), the Fondation pour la Recherche Médicale (EQU202303016295 to P.S.), the LabExs (ANR-10-LABX-73, REVIVE; ANR-10-LABX-0061, DEVweCAN; ANR-11-LABX-0042, CORTEX), and the University of Lyon within the program “Investissements d’Avenir” (ANR-11-IDEX-0007).

## Author contributions

Investigation: V.G., F.P., Y.P., N.D., N.A., C.C., I.A., P-Y.B.

Formal analysis: P-Y.B., P.S.

Writing original draft, Funding acquisition: P-Y.B., P.S.

Conceptualization, Supervision, Validation, Visualization, Project administration, Manuscript review and editing: P-Y.B., P.S.

---

**Supplementary figure 1:**
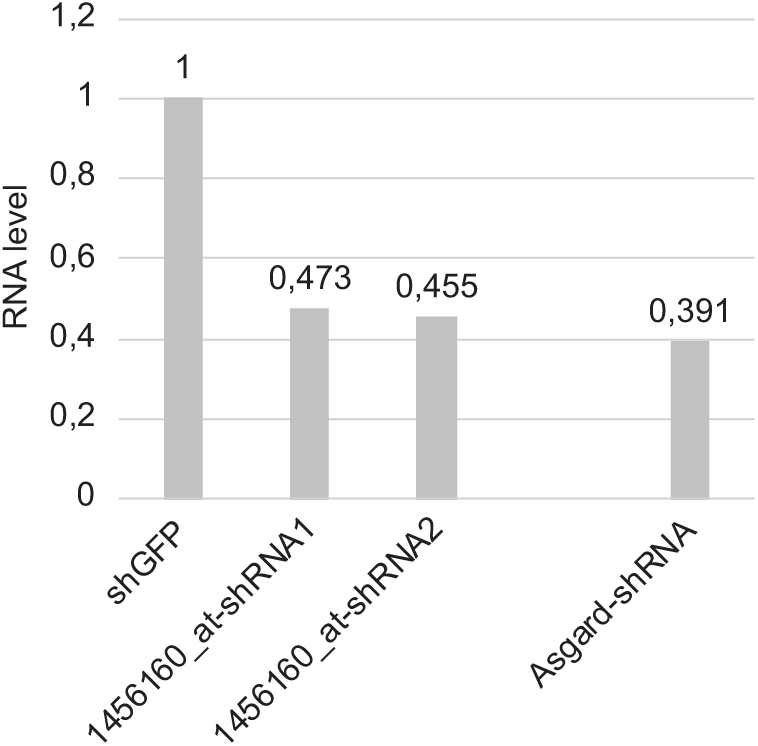
Validation of shRNA-mediated knockdown efficiency. qRT-PCR analysis of target gene expression following lentiviral transduction of mESCs with *1456160_at-shRNA1*, *1456160_at-shRNA2*, and *Asgard-shRNA*. Expression levels were normalized to those of cells transduced with a non-targeting control shRNA against GFP (shRNA-GFP).

**Supplementary Figure 2:**
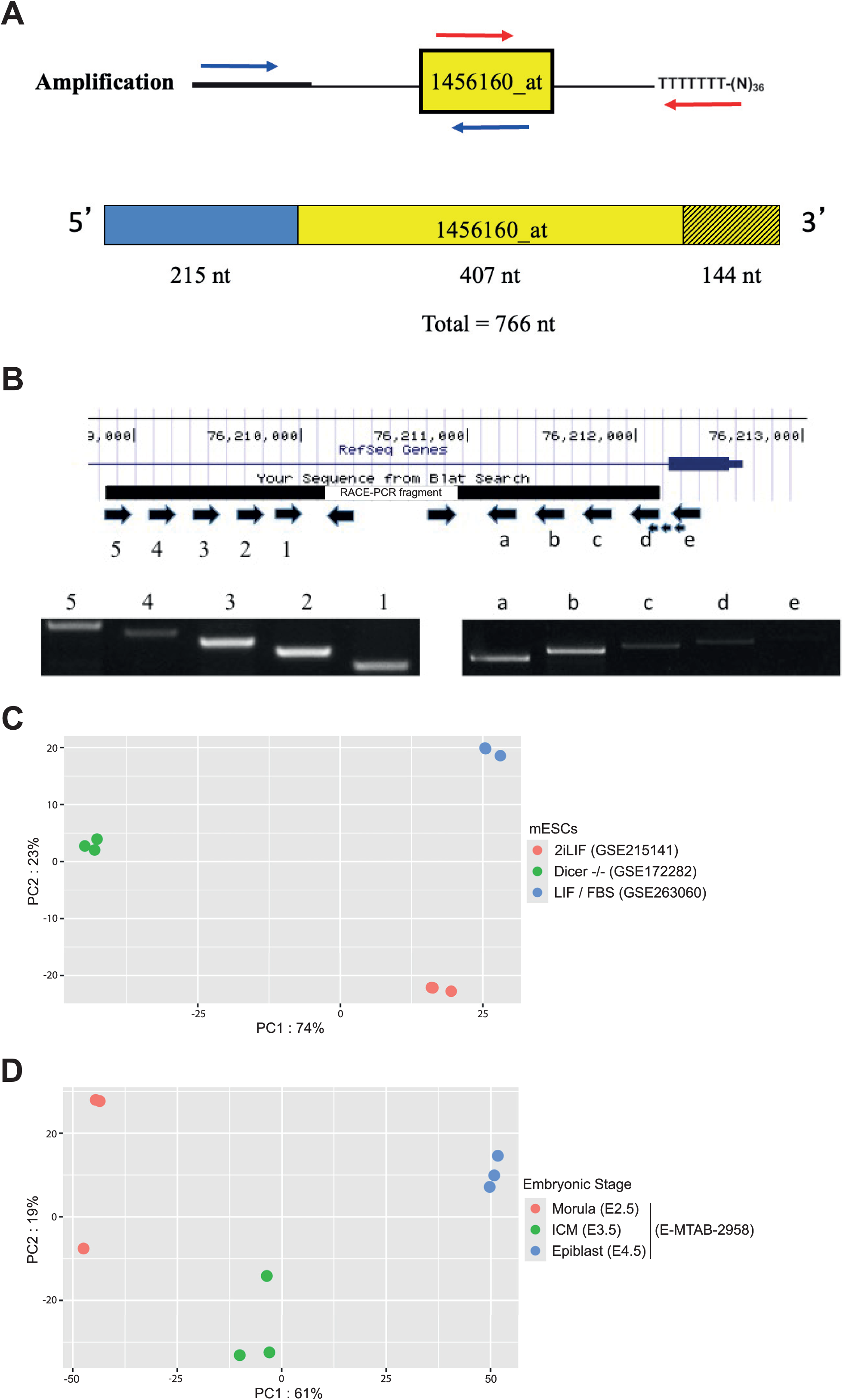
Experimental strategy for amplification of the full-length *Asgard* transcript. **(A)** Schematic representation of the RACE-PCR approach used to amplify a 766-nt fragment corresponding to the *1456160_at* sequence. **(B)** RT-PCR-based strategy for obtaining the full-length *Asgard* transcript using 5′ and 3′ primers located outside the RACE-amplified region. **(C)** Principal component analysis (PCA) of transcriptomes from E14tg2a mESCs cultured in LIF/serum or 2i/LIF conditions, and *Dicer^-/-^* mESCs, using datasets from GSE263060, GSE215141, and GSE172282. **(D)** Principal component analysis (PCA) of transcriptomes from mouse morula (E2.5), inner cell mass (ICM; E3.5), and epiblast (E4.5) stages, using data from E-MTAB-2958.

**Supplementary Figure 3:**
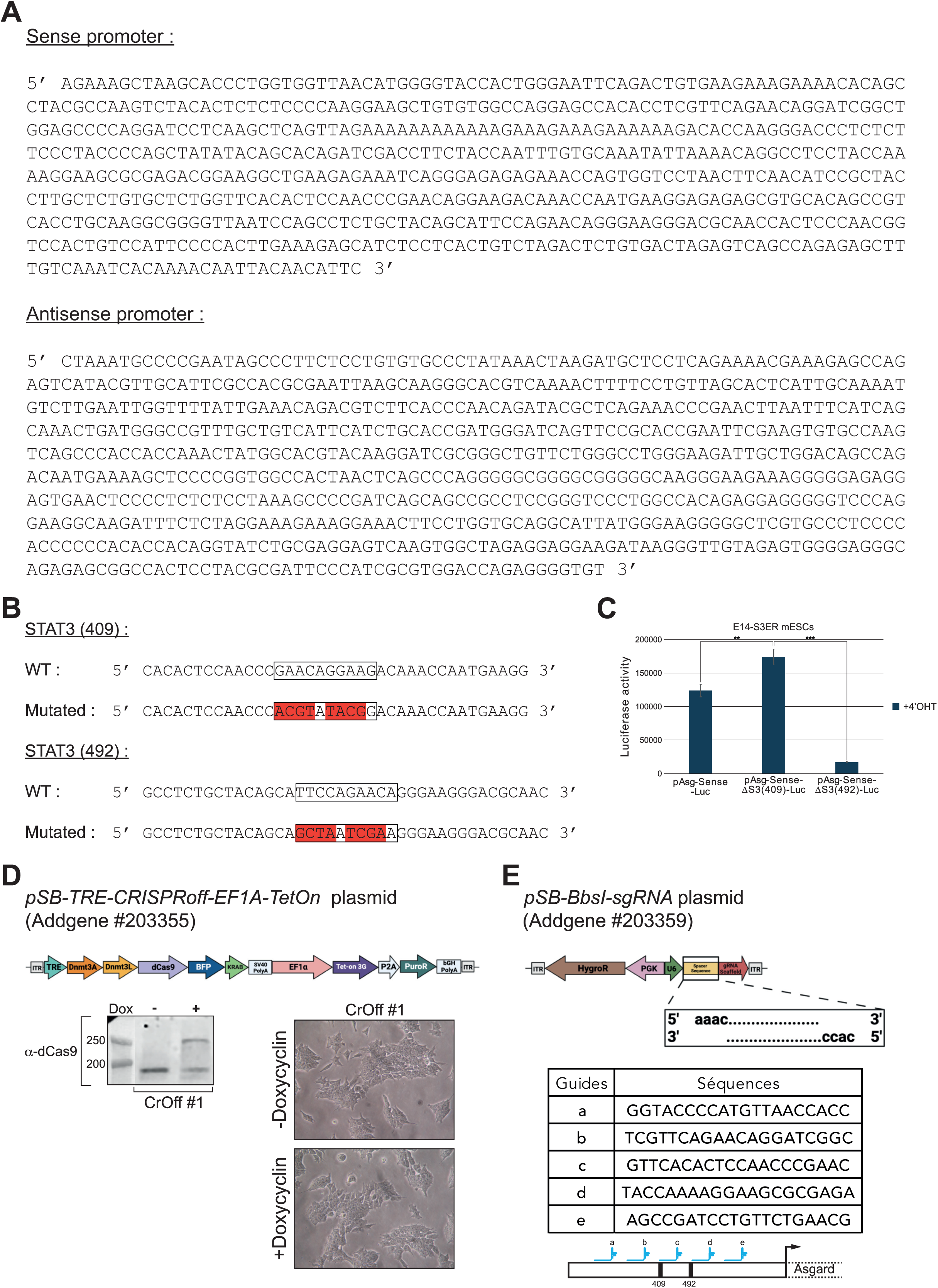
Functional analysis of STAT3 binding sites within the *Asgard* Sense promoter. **(A)** Nucleotide sequences of the “Sense” and “Antisense” promoter regions cloned into *pGL4-Luc* to generate the reporter constructs *pAsg-Sense-Luc* and *pAsg-Antisense-Luc.* **(B)** Sequences of the two putative STAT3 binding sites identified in the “Sense” promoter at positions 409 and 492, with mutations (in red) introduced into *pAsg-Sense-Luc* to generate the mutant constructs *pAsg-Sense-ΔS3(409)-Luc* and *pAsg-Sense-ΔS3(492)-Luc*. **(C)** Luciferase reporter assay in E14-S3ER mESCs transfected with either the wild-type or mutant constructs and cultured in the presence of 4’OHT. Data represent mean ± SE from three independent experiments. **(D)** Schematic representation of the *pSB-TRE-CRISPRoff-EF1A-TetOn* plasmid (Addgene #203355), enabling doxycycline-inducible expression of dCas9 fused to the DNMT3A and DNMT3L methyltransferases, as well as to the KRAB repression domain. Western blot analysis of dCas9 protein expression in the CrOff#1 clone of E14tg2a mESCs generated following transfection with the *pSB-TRE-CRISPRoff-EF1A-TetOn* plasmid and puromycin selection, cultured under –doxycycline and +doxycycline conditions. Phase-contrast images of CrOff#1 cells under –doxycycline and +doxycycline conditions. **(E)** Schematic representation of the *pSB-BbsI-sgRNA* plasmid (Addgene #203359), enabling expression of guide RNAs (sgRNAs). Table listing the sequences of five sgRNA spacer regions designed to target the *Asgard* gene promoter (designed using CRISPOR; https://crispor.gi.ucsc.edu). These sequences were cloned into the *pSB-BbsI-sgRNA* plasmid. Statistical significance was determined by Student’s *t*-test (\**p* < 0.05; \*\**p* < 0.01; \*\*\**p* < 0.001).

**Supplementary Figure 4:**
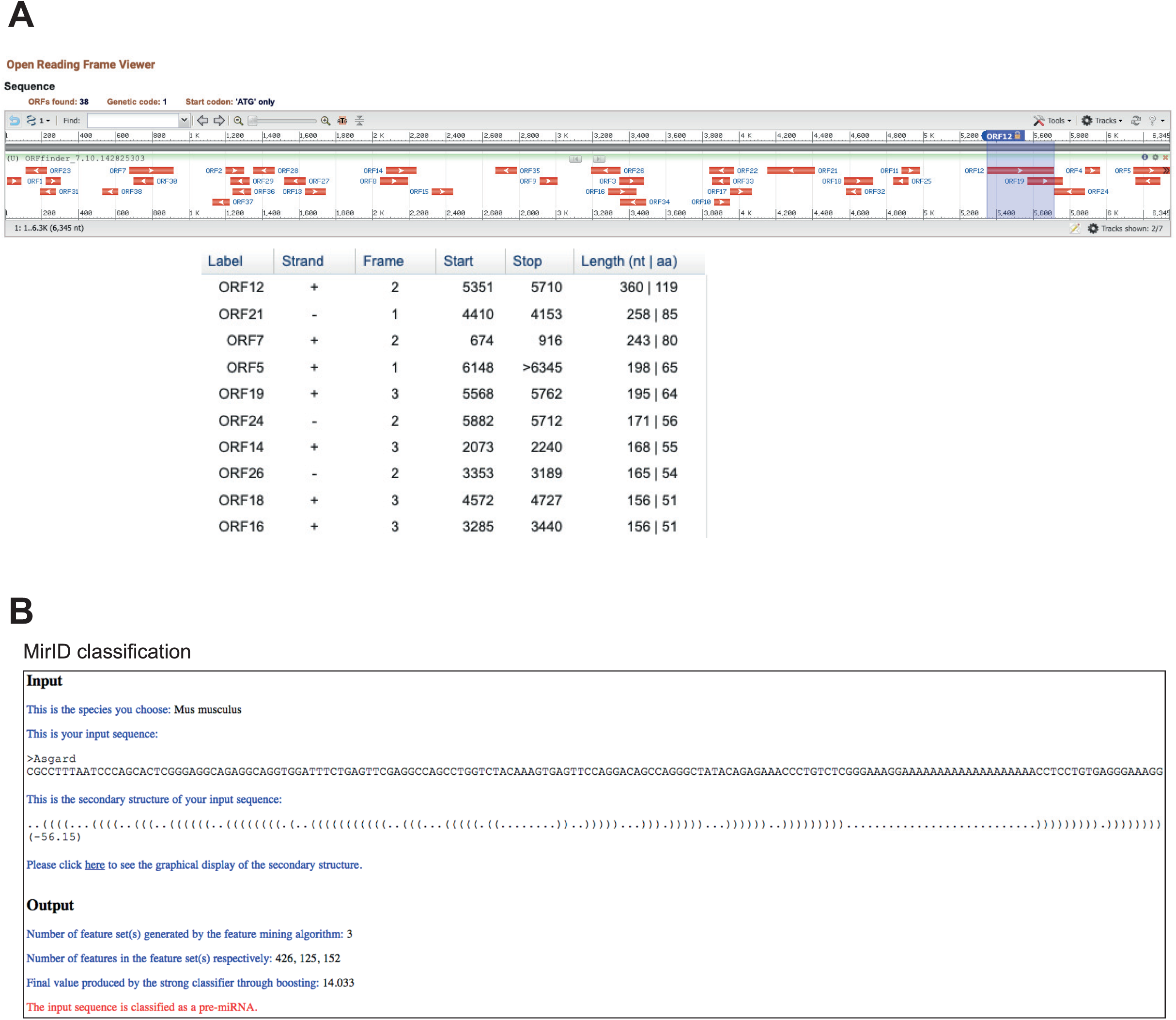
*In silico* analysis of the coding potential and miRNA biogenesis capacity of the *Asgard* transcript. **(A)** Open reading frame (ORF) prediction using the NCBI ORFfinder tool (https://www.ncbi.nlm.nih.gov/orffinder/). A total of 38 ORFs were identified, the longest spanning 360 nucleotides. **(B)** Computational prediction of the *Asgard* pre-miRNA sequence using the MirID algorithm (http://datalab.njit.edu/RNAcenter/). The input sequence was classified as a putative pre-miRNA, supporting its potential role in miRNA biogenesis.

**Supplementary Figure 5:**
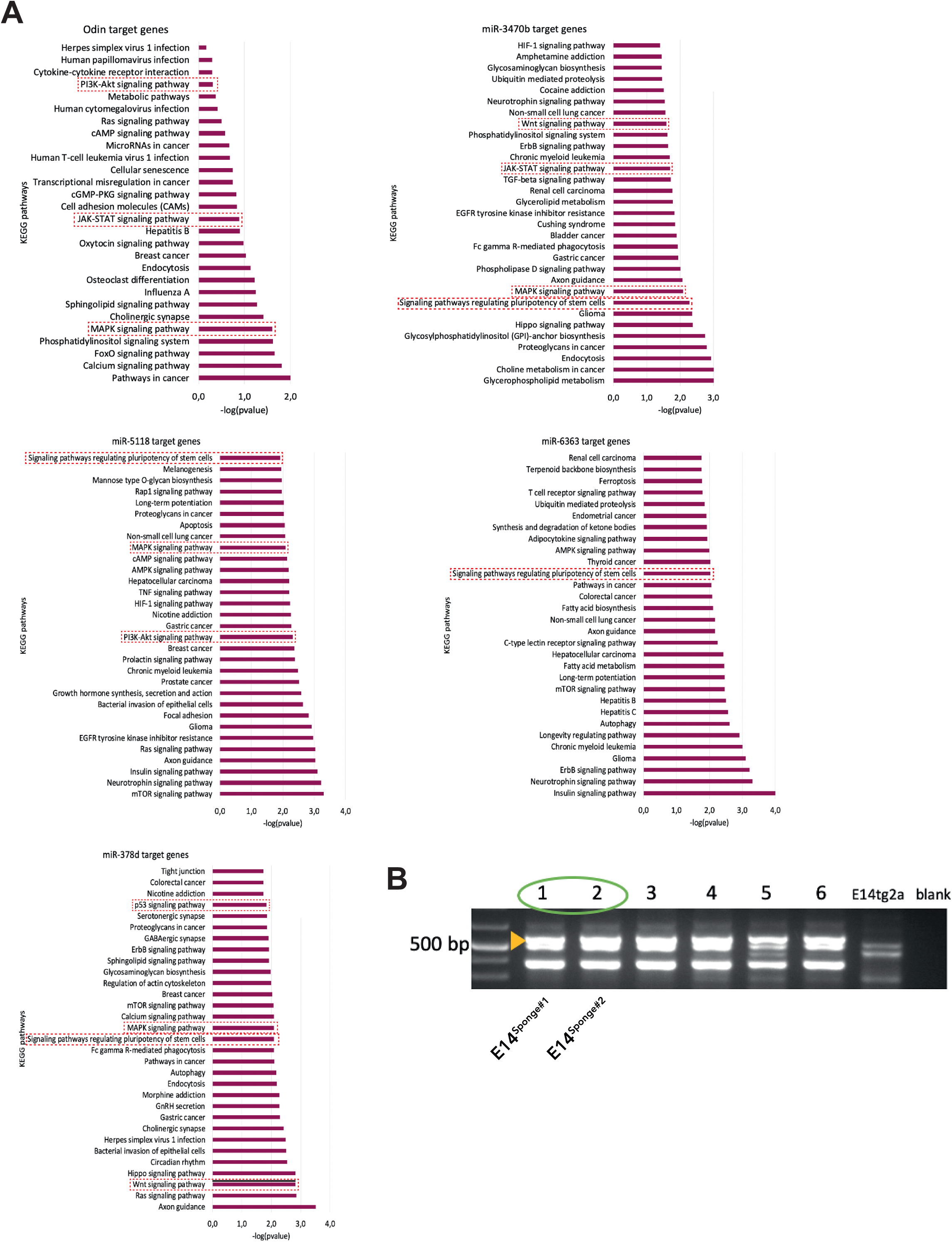
Functional pathway analysis of predicted miRNA targets and validation of *Asgard* sponge-expressing clones. **(A)** KEGG pathway enrichment analysis of predicted target genes for *Odin*, *miR-3470b*, *miR-5118*, *miR-6363*, and *miR-378d*, performed using the miRWalk platform. *P*-values were calculated using Fisher’s exact test to assess significant enrichment of KEGG pathways among the predicted targets. **(B)** PCR validation of six E14tg2a mESC clones generated by electroporation with the *pPB-Sponge* plasmid, followed by hygromycin selection and clonal expansion. Clones #1 and #2 were selected for further experiments and designated E14^Sponge#1^ and E14^Sponge#2^, respectively.

